# Characterizing Post-Mortem Brain Molecular Taxonomy of Cognitive Resilience and Translating it to Living Humans

**DOI:** 10.1101/2025.08.13.670106

**Authors:** Caio M. P. F. Batalha, Lei Yu, Andrea R. Zammit, Victoria N. Poole, Aron S. Buchman, Katia de Paiva Lopes, Ricardo Vialle, Peter Abadir, Lolita Nidadavolu, Tony Wyss-Coray, Nick T. Seyfried, Yanling Wang, Shinya Tasaki, Philip L. De Jager, Yasser Iturria-Medina, David A. Bennett

## Abstract

Here, we define cognitive resilience as slower or faster cognitive decline after we regress out the effects of common brain neuropathologies. Its understanding could provide important insights into the biology underlying cognitive health, enabling the development of more effective strategies to prevent cognitive decline and dementia. However, this requires the development of a practical method to quantify resilience and measure it in living individuals, as well as identifying heterogenous pathways associated with resilience in different individuals. Here, we approach this problem by using a data-driven framework to quantify and characterize molecular signatures underlying cognitive resilience. Using multimodal contrastive trajectory inference (mcTI) on bulk RNA sequencing and tandem mass tag (TMT) proteomic data from 898 post-mortem brain samples from the Religious Orders Study and the Rush Memory and Aging Project (ROSMAP), we derived individual-level molecular pseudotime values reflecting the molecular path from high to low resilience across individuals. Additionally, we identified two distinct molecular subtypes of resilience, each characterized by unique transcriptomic and proteomic signatures, and differing associations with several phenotypes. To translate our brain-derived pseudotime and subtypes to living individuals, we developed prediction models with paired genetics, ante-mortem blood omics, clinical, psychosocial, imaging and device data from the same individuals, demonstrating the potential to predict brain molecular resilience profiles in living persons. Our findings establish a framework for quantifying resilience based on multi- level molecular signatures, identify molecularly distinct resilience subtypes, and demonstrate the feasibility of translating brain-derived molecular profiles to living individuals—laying the groundwork for the development of targeted resilience-promoting interventions in cognitive aging.

## INTRODUCTION

Cognitive resilience refers to individuals that display less, or more, cognitive impairment than expected given their levels of brain pathologies (*1–3*). Some individuals show no sign of cognitive impairment, despite substantial brain pathologies, while some have more impairment relative to others with the same brain pathologies (*4–6*). This demonstrates substantial inter-individual differences in a brain’s ability to withstand injury and pathology. Thus, there are factors other than brain pathologies that contribute to or retard the rate of cognitive decline (*6–8*).

The current study extends our prior work that constructed and validated an operational definition of cognitive resilience (*9*). Previous work from our group has demonstrated the heterogeneity of pathologies in older brains with about 250 different combinations of brain pathologies (*10–12*). Yet, they only explain about 40% of the variance in cognitive decline and 2/3rds of Alzheimer’s dementia (*13*). By regressing out the effects of postmortem pathologic indices from longitudinal assessments of cognitive decline, we obtained a metric of residual cognitive decline (i.e. cognitive decline unrelated to pathologies), with some persons declining slower than predicted and others faster. We subsequently identified many multi-omic drivers associated with more or less resilience (*14–19*). Developing strategies to target resilience factors could prove to be a more cost-effective strategy than targeting each type of pathology individually as any intervention targeting resilience would offset any and all combinations of brain pathologies.

Advances in molecular data generation with high-throughput technologies for different omics layers have enabled the development of new methods capable of leveraging large amounts of data to untangle molecular signatures that may identify disease progression and heterogeneity. One approach, trajectory inference, also known as pseudotemporal ordering, enables the reconstruction of pseudolongitudinal trajectories of biological processes based on cross-sectional snapshots of omics data (*20–23*). These reconstructed trajectories are essentially a relative ordering of samples in relation to progression from a desired state to an undersired state according to their omics data patterns. We recently applied an advanced trajectory inference Machine-Learning (ML) method to multi-omic data from *post-mortem* brain samples to quantify person-specific pseudotime values that represent the extent of progression of participants along a continuum from a state of no cognitive impairment (NCI) to Alzheimer’s Disease (AD) dementia, and subsequently identified molecular subtrajectories, i.e., subtypes, that represent different molecular pathways from NCI to AD dementia (*24, 25*). The approach follows precision medicine approaches in cancer to identify molecular subtypes of disease which respond to different therapeutics despite identical clinical and pathologic features. Here, we apply the method to the molecular characterization of cognitive resilience to identify the molecular subtypes that progress from no residual cognitive decline to fast residual cognitive decline.

Specifically, we leveraged two sets of brain multi-omics data - bulk RNA sequencing and tandem mass tag (TMT) proteomics – and integrated their molecular information through the advanced ML algorithm (*24, 25*). Additionally, we leveraged ante-mortem data from the same individuals to develop prediction models for pseudotime and for the subtypes, as a step to translating the molecular pseudotime and subtypes to living persons.

## RESULTS

### Analytical approach

We used data from individuals in the Religious Orders Study or Rush Memory and Aging Project (ROSMAP) cohorts (see Methods). We started by defining the desired (background) and undesired (target) states using a latent class functional mixed-effects model to cluster ROSMAP participants (N = 1409) based on their patterns of residual cognitive decline (*26*). This resulted in four clusters displaying distinct patterns of decline, with resilience cluster 1 representing individuals with no residual cognitive decline (more resilience), resilience cluster 4 representing individuals with the most rapid residual cognitive decline (less resilience), and resilience clusters 2 and 3 with intermediate levels of decline (Figure 1A). Cluster 1 was used as the background, and clusters 3 and 4 were merged and used as the target, as cluster 4 had too few persons for a stable model.

**Figure 1.**
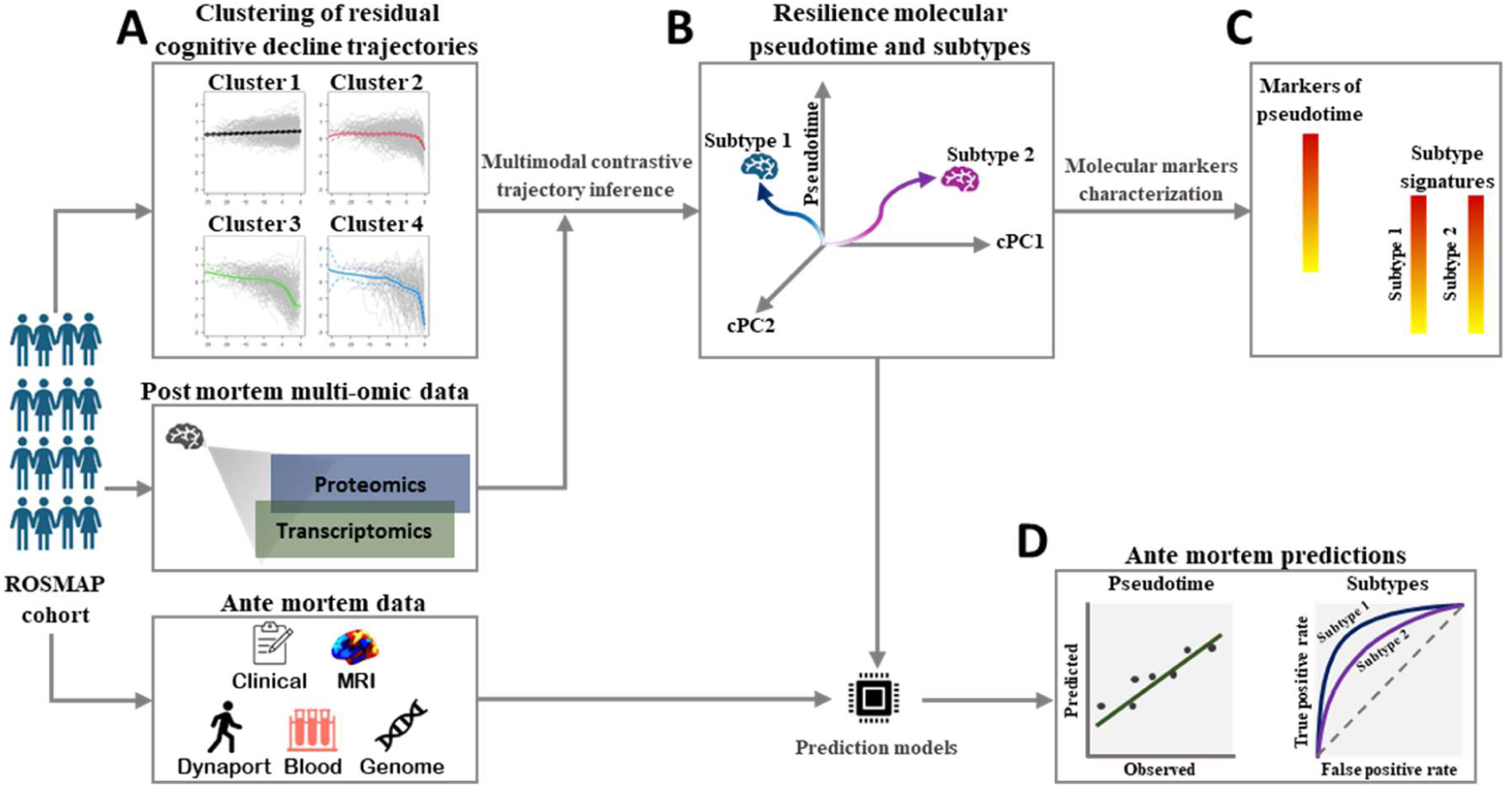
Overview of our analytic approach. A) Residual cognitive trajectories of ROSMAP participants (i.e., after neuropathologies adjustment) are clustered by their trajectories and used to define a background (cluster 1) and a target (clusters 3 and 4) dataset. B) *Post-mortem* transcriptomic and proteomic data from the same individuals are then integrated via a contrastive trajectory inference analysis, resulting in a molecular pseudotime estimation, and distinct subtypes that account for different molecular pathways from high to low resilience. C) Molecular features associated with pseudotime and subtypes are then characterized. D) Finally, *ante- mortem* multi-modal data from the same individuals is used to build classifiers to translate the brain molecular pseudotime and subtypes to living participants.

For analyses, we used all participants with a resilience cluster classification and dorsolateral prefrontal cortex bulk RNA-Seq (n=844) and/or tandem mass tag (TMT) proteomics (n=613) for a total n=898. The omics data was analyzed using a machine learning (ML) algorithm, mcTI - multimodal contrastive trajectory inference (*24, 25*), that characterizes the molecular patterns in the target phenotype of interest – here, less resilience. The main outputs are (i) a person-specific pseudotime value and (ii) the clustering of each individual into a mutually exclusive subtype.

Pseudotime is a unidimensional value that reflects the severity of residual cognitive decline (i.e., resilience), with values closer to zero indicating more resilience and values closer to one indicating less resilience. Therefore, positive associations with pseudotime are related to faster residual cognitive decline. Conversely, negative associations with pseudotime are related to slower residual cognitive decline. Subtypes are clusters of individuals based on the molecular patterns in the omics data revealed by mcTI. They represent individuals who, despite having similar pseudotime values, have distinct molecular pathways associated with their resilience phenotype. In essence, subtypes represent different subtrajectories from high to low resilience. The clustering procedure is carried out in a lower dimensional space constructed via t-SNE embedding (*27*). The optimum number of subtypes is chosen using a majority rule across the Calinski–Harabasz and Silhouette criteria (*28, 29*), and the stability and significance of subtypes are evaluated via randomized permutations (see Methods for a more detailed description).

Figure 1 shows a general schematic of our analyses, including downstream steps of characterization of markers of resilience molecular pseudotime and subtypes, and the analysis of the ability of *ante-mortem* data from the same individuals to predict these two brain-derived variables.

### Estimation of molecular pseudotime reflecting brain residual cognitive decline severity

As expected, molecular pseudotime reflects the severity of resilience clusters (Figure 2A). Also as expected, pseudotime displayed strong negative correlations with the rate of change of multiple cognitive domains assessed longitudinally prior to death (all FDR-adjusted P<0.0001), including residual cognitive domains associated with our operational definition of resilience (Figure 2B). Memory related cognitive domains (episodic memory, semantic memory and working memory) displayed the strongest correlations with resilience pseudotime. These results indicate that higher pseudotime values (lower resilience) are associated with faster cognitive decline (steeper negative slopes).

**Figure 2.**
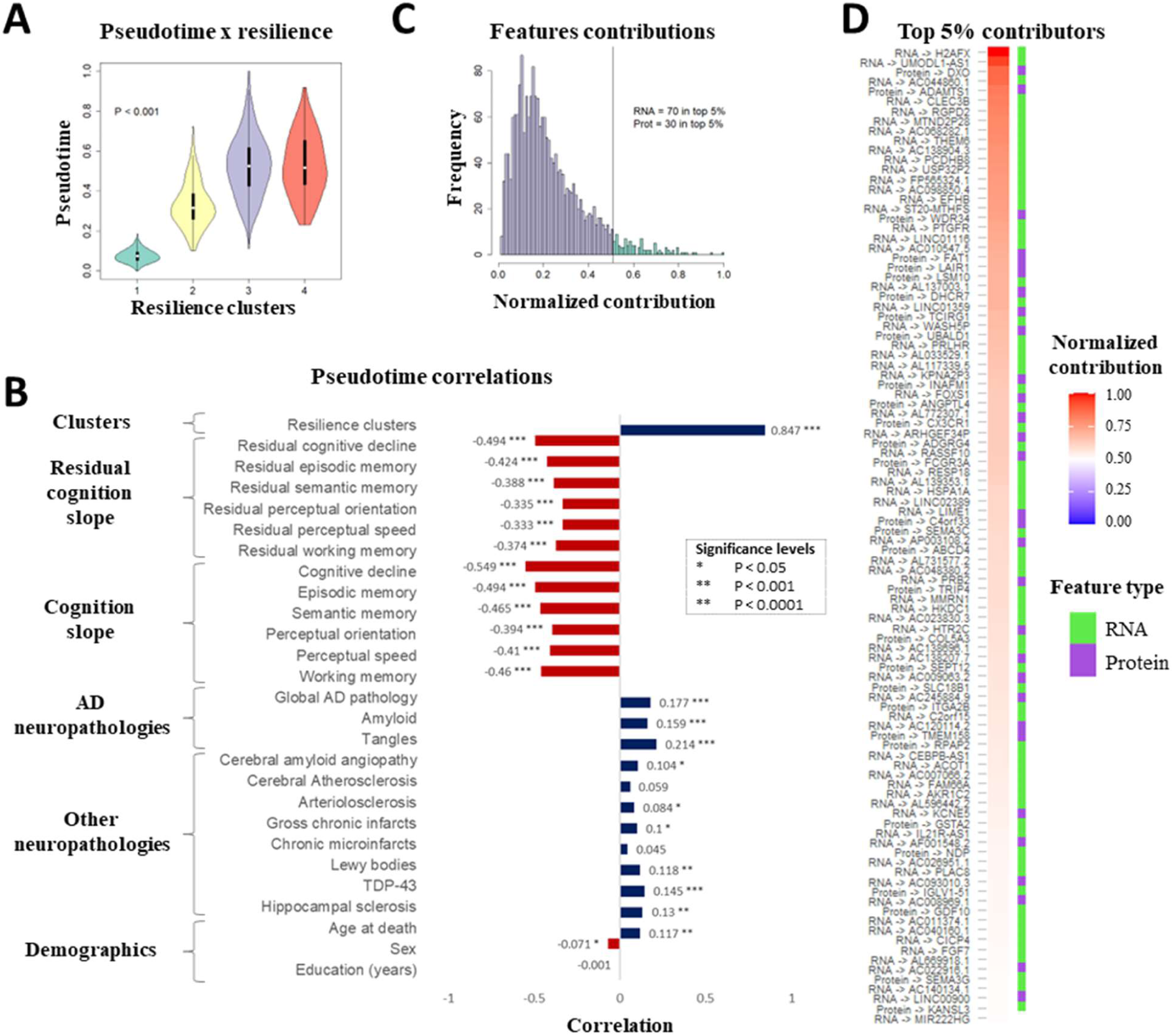
Estimation of molecular pseudotime reflecting brain residual cognitive decline severity. A) Resilience pseudotime association with resilience clusters, based on ANOVA with 5,000 permutations. B) Resilience pseudotime correlations with the rate of change of cognitive domains and their residual counterparts, neuropathologies, and demographic variables; positive (blue bars) and negative (red bars) correlations are displayed with varying levels of significance (*: FDR-adjusted P < 0.05; **: FDR-adjusted P < 0.001; ***: FDR-adjusted P < 0.0001). C) Distribution of features contributions to the construction of the molecular pseudotime variable; contributions were calculated based on the contrastive PC (cPC) weights of features and normalized by the highest contributor in both omic layers (see Methods); the green colored area represents the top 5% contributors to pseudotime. D) Top 5% contributors to pseudotime (see Methods - Assessing features’ contributions to pseudotime).

Although the resilience clusters were derived from a model that controlled for demographics and neuropathologies, pseudotime had modest associations with age and sex, and AD neuropathologies and other pathologies (FDR-adjusted P<0.05), the strongest for global AD pathologic burden, amyloid-β load, tangle density and TDP-43 staining. (Figure 2B). These residual associations were observed in prior work (*18, 30*). This likely reflects some molecular interaction between these neuropathologies and residual cognitive decline that cannot be disentangled, suggesting that some features have pleiotropic effects related to these variables. There were no associations with cerebral atherosclerosis and chronic microinfarts.

We next quantified the contributions of each molecular feature to the resilience pseudotime based on the loadings of the cPC (contrastive principal component) matrix calculated by the mcTI algorithm (see Methods). The distribution of features shows that the contributions are highly unequal, with few features driving most of the pseudotime value (Figure 2C), and a larger proportion of features from the transcriptome among the top 5% contributors.

Figure 2D shows the top contributors to resilience molecular pseudotime. The top contributor is the transcript of *H2AFX*, a variant of histone *H2A*, one of the four core histones that compose the nucleosome, and is responsible for regulating DNA accessibility (*31*).

### Biological processes associated with resilience molecular pseudotime

While the top contributors to pseudotime are determined based on the 1000 mRNA and 1000 proteins preselected for the trajectory inference (see Methods), these preselected features are a small subset of the total number of features in the data, and they are not likely to be the only features associated with resilience. To better understand this global picture of the biological pathways associated with resilience pseudotime, we tested the association of pseudotime with all features in the dataset, and did a GO enrichment analysis of the significant results.

We first identified features significantly associated with the pseudotime through linear regression (Figure 3A). 2,987 RNA features and 756 protein features were positively associated with pseudotime, and 3,292 RNA features and 1,136 protein features were negatively associated with pseudotime (FDR-adjusted P < 0.05). Out of these results, 282 RNA features (154 positively and 128 negatively associated with pseudotime) were among the 1000 preselected RNA features for mcTI. For protein features, 127 (97 positively and 30 negatively associated with pseudotime) were among the 1000 preselected protein features for mcTI. This means that most features associated with pseudotime were not used as inputs for mcTI, highlighting the need to analyze these additional features to have a better understanding of the biological processes underlying resilience. The full regression results are available in File S1. We then selected the top 500 based on R^2^ in each category (positive RNAs, negative RNAs, positive proteins and negative proteins), and performed a GO enrichment analysis.

**Figure 3.**
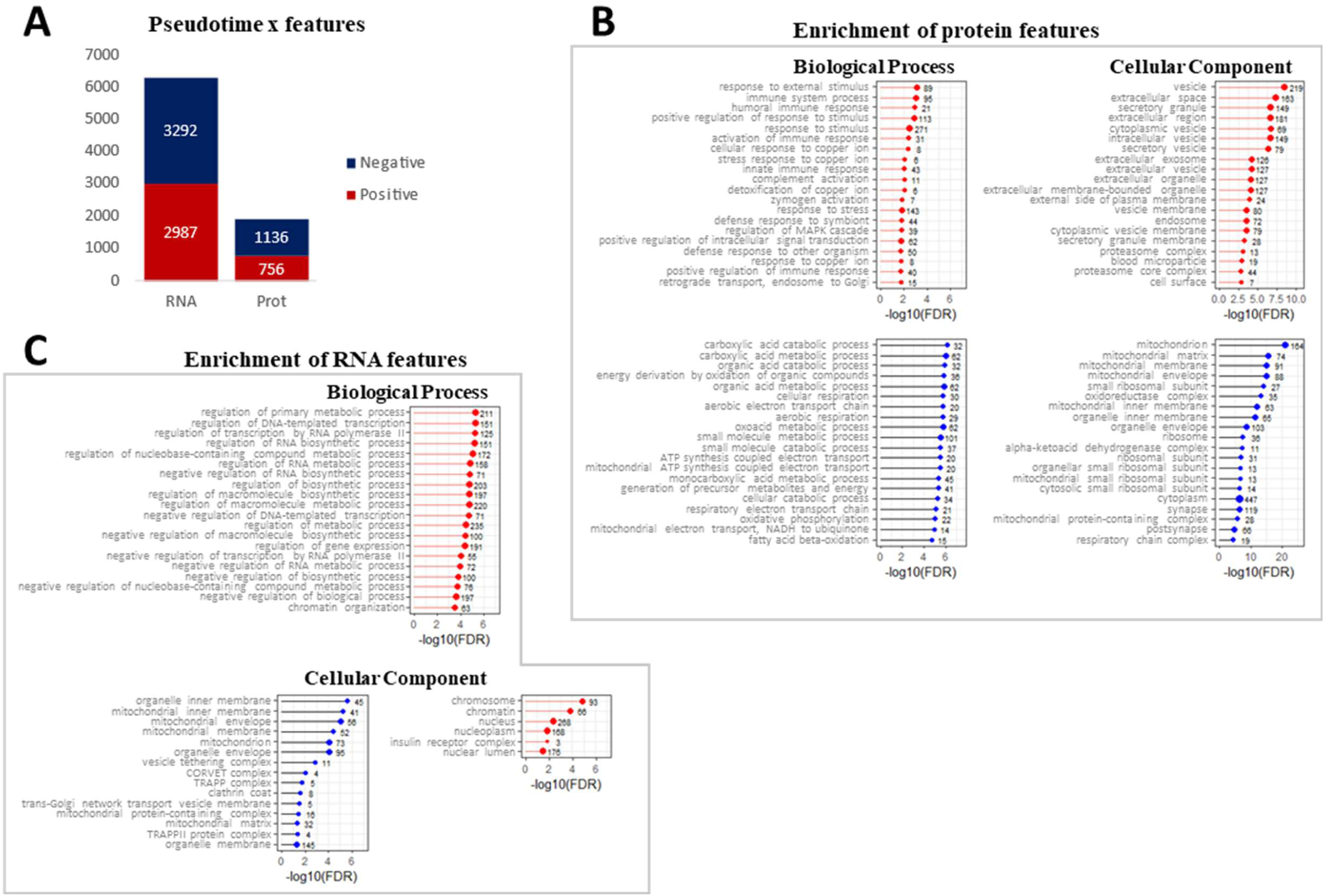
Biological processes associated with resilience molecular pseudotime. A) Molecular features associated with resilience pseudotime, based on linear regression; red and blue portions of each bar represent, respectively, features positively and negatively associated with pseudotime (all with FDR-adjusted P<0.05). B & C) GO enrichment analysis of the top 500 positively (red colored charts) and top 500 negatively (blue colored charts) associated RNA (C) and protein (B) features with pseudotime (FDR-adjusted P<0.05), selected based on the highest R^2^ values in the linear regression; only the top 20 enriched terms per category are show in the charts (the full list of enriched terms can be found in File S2); numbers next to each bar in the plot represent the number of features (among the 500 used) annotated to that term.

Results show that features negatively associated with pseudotime display a common enrichment pattern in mitochondria-related terms (Figures 3B and C, blue colored charts). For protein features, this was found both for Biological Process and Cellular Component terms (Figure 3B). For RNA features, there were no enrichments for Biological Process, but Cellular Component revealed enrichment in mitochondrion and mitochondrial sub-compartments (Figure 3C).

Together, these results show that the continuum of pseudotime from high to low resilience is accompanied by lower expression of features involved in mitochondrial processes and contained in the mitochondrial compartment.

Features positively associated with pseudotime displayed more variation (Figures 3B and C, red colored charts). For RNA, enriched terms centered around regulation of gene expression, with the nuclear compartment as cellular component (Figure 3C). For proteins, there was a prominence of immune system related processes, with extra cellular regions as cellular component (Figure 3B).

#### Characterization of resilience molecular subtypes

Next, we identified different subtrajectories of pseudotime progression that represent distinct molecular subtypes of resilience. These subtrajectories or subtypes are clusters of individuals that display distinct patterns in multi-dimensional contrastive principal component (cPC) space. The optimum number of subtypes was calculated based on a majority rule across several criteria (*28, 29*) (see Methods).

We identified two distinct resilience molecular subtrajectories along the pseudotime axis (see Figure 4A, for a representation of pseudotime subtrajectories in two-dimensional cPC space). Out of the 488 non-background samples, 212 (∼43%) were classified as subtype 1 and 276 (∼57%) were classified as subtype 2 (Figure 4B).

**Figure 4.**
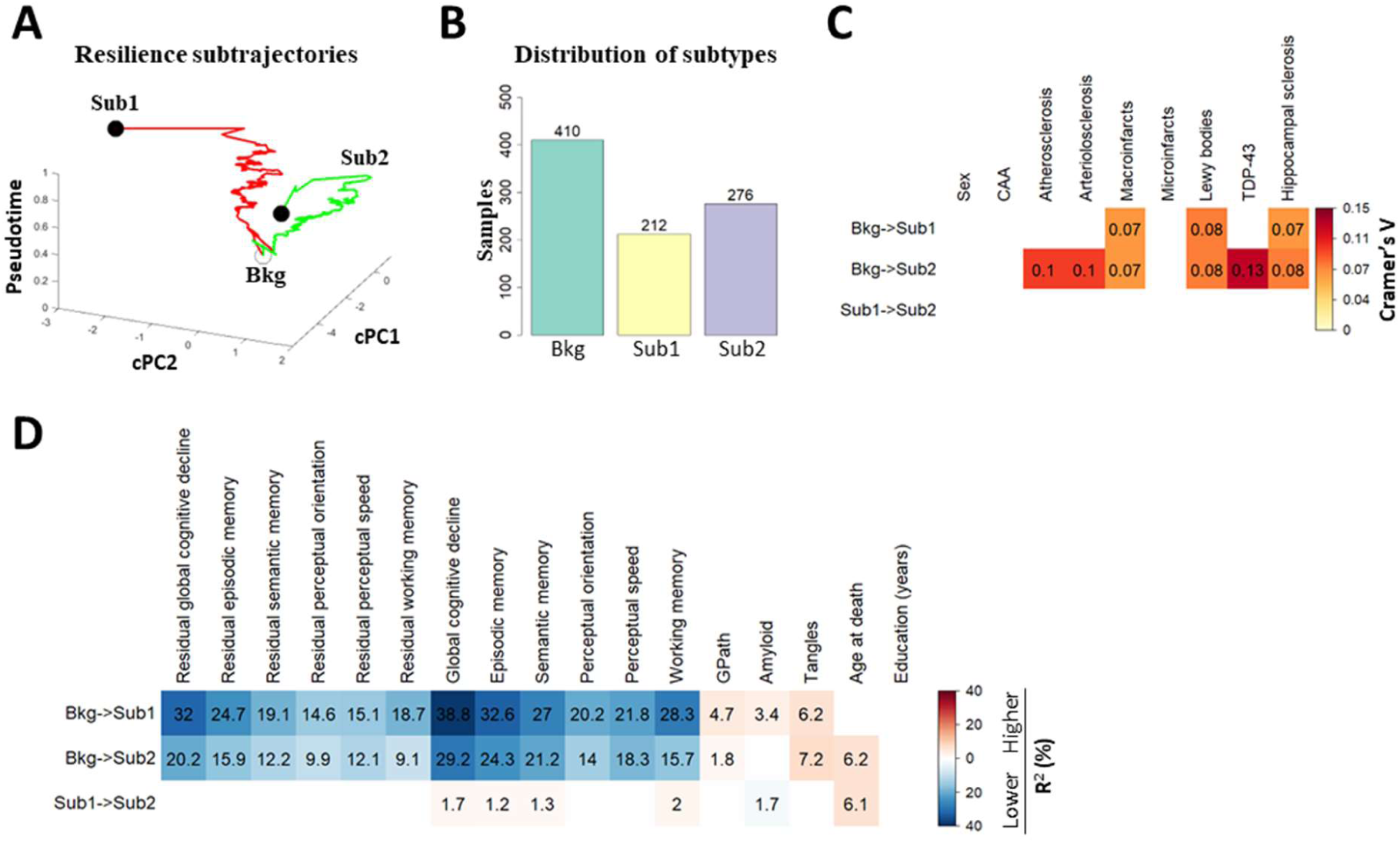
Characterization of resilience molecular subtypes. A) Visualization of two pseudotime subtrajectories in relation to the two first cPCs. Both subtypes contain individuals who progress along the pseudotime axis from the least severe cases (near background samples – Bkg) to the most severe cases (closed black circles), but they do so along different trajectories. B) Distribution of participants among the two identified subtypes and the background. C) Association of subtypes with different cognitive, pathologic and demographic variables – only categorical variables, tested through Chi-squared test. Numbers and cell shade intensities represent Cramer’s V strength of association. Blank cells had non-significant differences (FDR- adjusted P>=0.05). CAA for cerebral amyloid angiopathy. D) Association of subtypes with different cognitive, pathologic and demographic variables – only continuous variables, tested through ANOVA with 5,000 permutations. Red and blue shaded cells represent, respectively, positive and negative differences in the group comparisons (Bkg->Sub1, Bkg->Sub2 and Sub1->Sub2). Numbers inside cells represent R^2^ (%) values. Blank cells had non-significant differences (FDR-adjusted P>=0.05).

Different categorical variables were compared using a Chi-squared test (Figure 4C), while quantitative variables across subtypes were compared with permutation ANOVA (Figure 4D). Subtype 2 individuals were older relative to background and subtype 1, while subtype 1 individuals were similar in age to background individuals (Figure 4D; FDR-adjusted P < 0.05). There were no differences in sex or years of education (Figures 4C, D; FDR-adjusted P < 0.05). Both subtypes display more pronounced cognitive decline across all domains when compared to background samples (the most resilient individuals), as expected, but subtype 2 displayed less decline. When subtypes 1 and 2 are compared directly, there are differences in global cognitive decline, and the episodic memory, semantic memory and working memory domains (Figure 4D; FDR-adjusted P < 0.05).

A similar pattern is observed for a summary measure of AD pathology, with less pathology in subtype 2 vs subtype 1. Further, greater amyloid-β load could only be detected in subtype 1, while greater PHF-tau tangle density is found in both subtypes (Figure 4D; FDR-adjusted P < 0.05). For non-AD neuropathologies, when compared to background individuals, both subtype 1 and 2 displayed significantly higher levels of macroinfarcts, Lewy bodies and hippocampal sclerosis, while subtype 2 also displayed additional higher burdens of atherosclerosis, arteriolosclerosis and TDP-43 (Figure 4C; FDR-adjusted P < 0.05). These differences, however, are small, and not significant in the direct comparison between subtypes 1 and 2 (Figure 4C, last column).

#### Distinct molecular signatures of resilience subtypes

To characterize the molecular differences contributing to each resilience subtype, we identified distinct omic features, among preselected features used as inputs for mcTI, differentially expressed in one subtype (in relation to background samples) but not in the other (FDR-adjusted P < 0.05). These subtype-specific differentially altered features represent the molecular underpinnings of the distinct subtrajectories (see Methods - Identification of distinct subtype-specific signatures).

This resulted in 113 unique features characterizing resilience subtype 1 signature, and 372 unique features characterizing resilience subtype 2 (Figures 5A and B). Subtype 1 displayed a roughly similar distribution of RNA and protein features among its signature (45% and 55%, respectively; Figure 5B – intra-subtype proportions), while subtype 2 displayed a higher representation of RNA features, totaling nearly two thirds of the signature features (64% and 36%, respectively; Figure 5B – intra-subtype proportions). Subtype 2 had a higher number of unique features in both omic layers, with the 237 distinct RNA features representing 82% of the total uniquely altered RNA features in both subtypes, and the 135 protein samples representing 69% of the uniquely altered protein features in both subtypes (Figure 5B – intra-modality proportions). Figure 5C presents the top 30 distinct contributors for each subtype (a complete list of distinct contributors for each subtype can be found in File S3).

**Figure 5.**
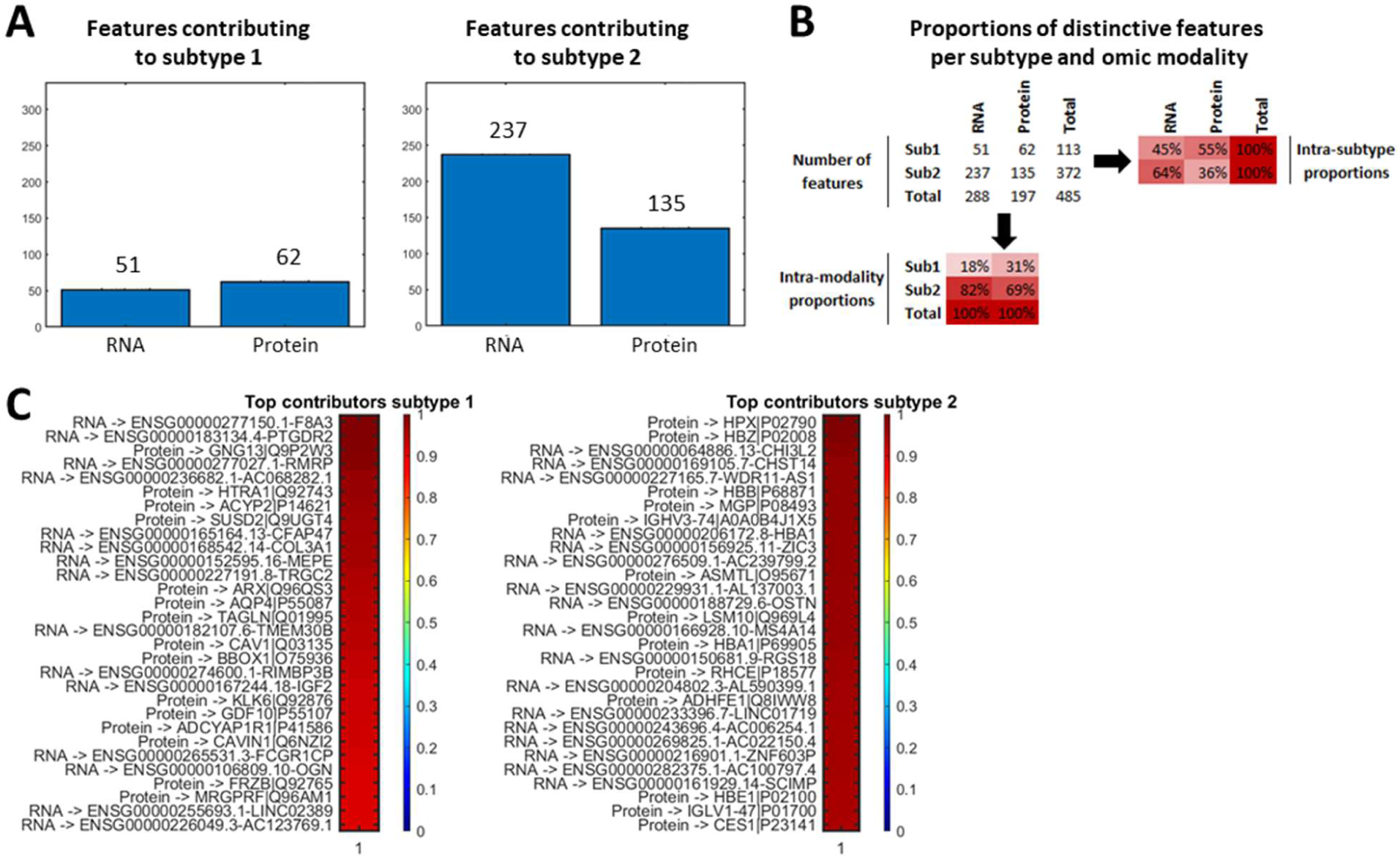
Distinct molecular signatures of resilience subtypes. A) Number of distinct features contributing to each subtype. B) Proportions of distinct features per subtype in relation to the overall number of altered features. C) Top 30 distinct molecular features altered in each subtype (full list in File S3).

#### Ante-mortem predictors of brain resilience molecular pseudotime

For brain molecular pseudotime and subtypes to ever be clinically actionable, it’s essential that they be translated to living humans. Here we leverage a wide range of data collected from the same persons prior to death. We previously translated molecular pseudotime and subtypes of AD to living individuals (*25, 32*). Here, we extend that work to resilience. We used blood molecular data (monocyte RNA, somalogic proteomics, metabolomics, cytokines, circulating cell-free DNA, pTau-217, pTau-181, routinary blood tests), psychosocial risk factors, sensor-derived mobility indices, MRI-derived measures, and polygenic scores (PGS). These data were used for two types of analyses: (i) association tests, and (ii) testing prediction models with cross-validation.

First, we tested the associations between blood transcriptomic, proteomic and metabolomic markers and pseudotime with linear regression controlling for age of death, sex and years of education. The partial R^2^ for each feature was obtained by comparing with the reduced regression model without features. A total of 33 blood markers were associated with brain resilience molecular pseudotime (FDR-adjusted P<0.05), most of which were metabolites with negative associations (Figure 6A), i.e., associated with progression from more to less resilience.

**Figure 6.**
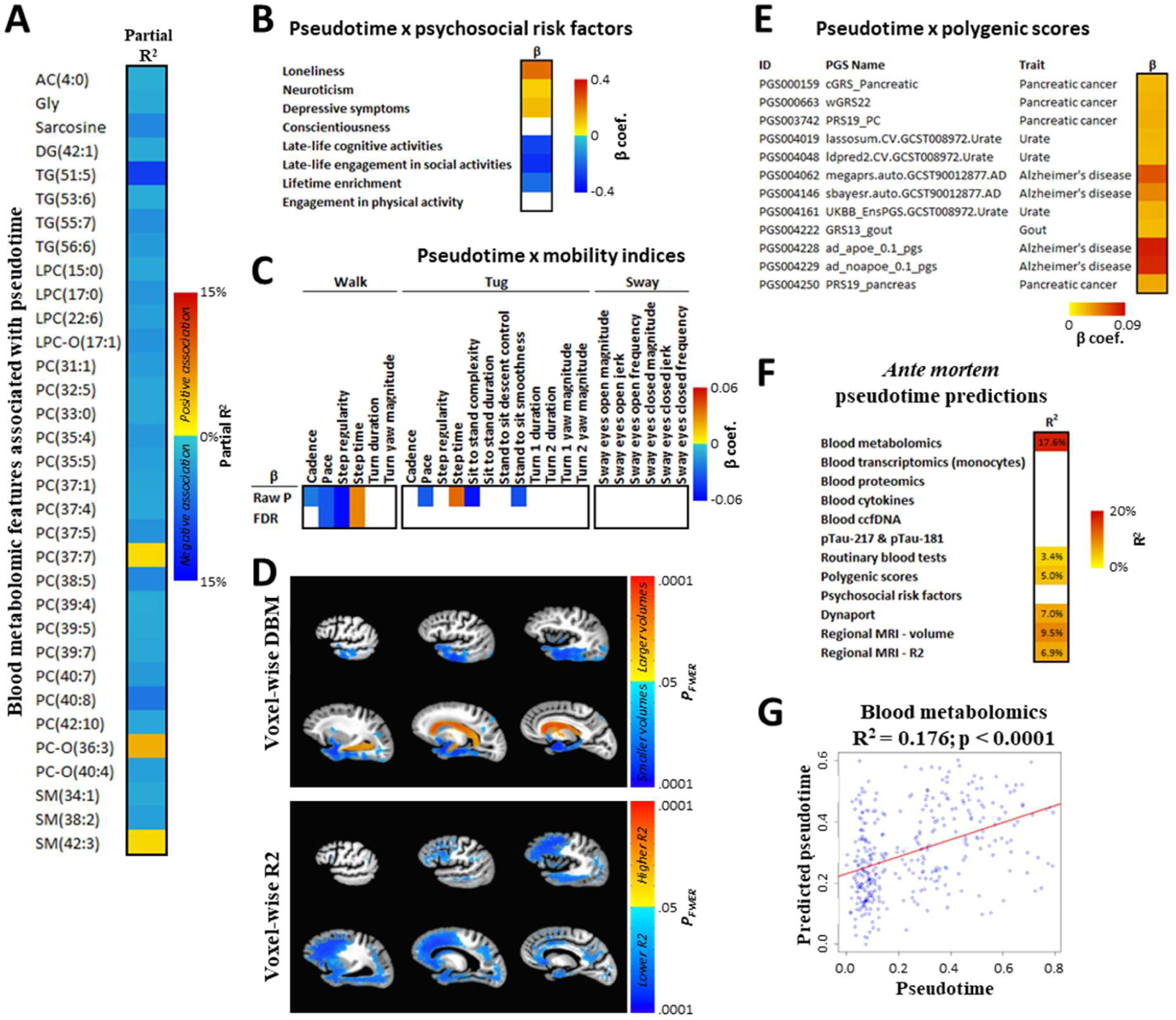
*Ante-mortem* predictions of *post-mortem* brain-derived molecular resilience pseudotime. A) Blood metabolomic markers significantly associated with brain-derived pseudotime (FDR-adjusted P<0.05). No blood transcriptomic or proteomic feature displayed a significant association. AC: acylcarnitines; DG: diglycerides; TG: triglycerides; LPC: lysophosphatidylcholines; PC: phosphatidylcholines; SM: sphingomyelins. B) Association of psychosocial risk factors with brain-derived resilience pseudotime. Red and blue shaded cells represent, respectively, positive and negative beta coefficients significantly associated with pseudotime (P<0.001). Blank cells had non-significant associations (P>=0.001). C) Association of mobility indices with brain-derived resilience pseudotime. Red and blue shaded cells represent, respectively, positive and negative beta coefficients significantly associated with pseudotime. Both raw p-values and FDR-adjusted p-values using M_eff_ are presented (both considered significant at <0.05). Blank cells had non-significant associations (FDR-adjusted P>=0.05). D) Associations of *post-mortem* brain structure with brain-derived molecular resilience pseudotime. Warm and cold colors indicate the magnitude of positive and negative associations of regional brain deformation (i.e., volumes) and R2 (i.e., tissue integrity), respectively (see Methods – Neuroimaging analysis). E) Associations of polygenic scores with brain-derived molecular resilience pseudotime. Red shade intensity represents beta coefficients. Only significant associations (FDR-adjusted P<0.05) are shown. F) *Ante-mortem* cross-validated predictions of brain-derived molecular resilience pseudotimes. Red shade intensity and numbers inside cells represent R^2^ (%) values of significant predictions (P<0.05). Blank cells had non-significant predictions (P>=0.05). G) Scatterplot of observed pseudotime vs predicted pseudotime for the best performing prediction, made with blood metabolomic data.

Next, we assessed whether psychosocial risk factors were associated with resilience molecular pseudotime. Pseudotime associations were tested separately for: loneliness, neuroticism, depressive symptoms, conscientiousness, late-life engagement in cognitive activities, late-life engagement in social activities, lifetime enrichment, and engagement in physical activity (Figure 6B). For psychological traits, higher pseudotime was positively associated with loneliness, neuroticism, and depressive symptoms (P < 0.001), as expected given their association with cognitive decline but not brain pathologies. No association was detected for conscientiousness. For lifestyle factors, higher pseudotime was inversely associated with engagement in late-life cognitive activities, late-life engagement in social activity, and lifetime enrichment (P < 0.001), as expected. No association was detected for engagement in physical activity.

Additionally, we assessed the association of quantitative sensor-derived gait and balance indices with molecular pseudotime using gait measurements collected with DynaPort (*33–35*), a belt-worn sensor with a triaxial accelerometer and three gyroscopes that captures acceleration and angular velocity in 3 directions. Gait measurements from the sensor were grouped into three performances: walk (32 ft), tug (timed up and go, composed of the subtasks: transition from sit to stand, transition from stand to sit, and turning) and sway (see Methods – DynaPort for more details on the variables). Since the variables composing these tasks have correlations with each other, doing a regular FDR adjustment for multiple tests could lead to an overcorrection. Because of this, we performed multiple test correction using the effective number (M_eff_) of independent tests (*36*), using the FDR implementation proposed by Ji & Li (2005) (*37*). Our results show a negative association of pseudotime with pace speed and step regularity, indicating reduced gait speed in individuals with less resilience (Figure 6C). Step time was increased, reinforcing evidence for slower gait speed. Given that this analysis has a relatively limited sample size (N = 185), we present significant raw p-values, which could represent potential true associations.

Negative associations with raw p-values included decreased walk cadence, as well as pace, sit to stand complexity and stand to sit smoothness. Step time in TUG tasks was also positively associated with pseudotime considering raw p-values.

To characterize brain structural associations with resilience pseudotime, we analyzed *post-mortem* deformation-based morphometry (DBM) and R_2_ data due to limited *ante-mortem* imaging data in this sample. Our results show that smaller temporal regions and larger ventricles were significantly associated with higher pseudotime values (FWER-adjusted P<0.05) (Figure 6D) (see Methods - Neuroimaging analysis).

We also assessed pseudotime associations with 802 polygenic scores, and our results reveal 12 PGS significantly associated with pseudotime (FDR-adjusted P<0.05), all with a positive association (Figure 6E). These PGS are associated with Alzheimer’s disease (PGS004062, PGS004146, PGS004228, PGS004229), serum urate levels (PGS004019, PGS004048, PGS004161), gout (PGS004222), and pancreatic cancer (PGS000159, PGS000663, PGS003742, PGS004250). Urate is the predominant form of uric acid at physiologic pH (*38*). Increased urate concentrations are associated with the deposition of monosodium urate crystals, causing the development of gout (*39*), and previous studies have shown an association of uric acid with cognitive function (*40, 41*). Cognitive impairment has been observed in different types of cancer (*42*), although it’s unclear to us why we found associations only for PGS related to pancreatic cancer.

Finally, we evaluated the predictive power of data modalities to predict resilience molecular pseudotime by using several prediction algorithms with cross-validations (prediction algorithms are described in Methods – Statistical analyses). Each data modality was analyzed separately due to the low number of samples overlapping different modalities. Results show that the data modality that demonstrated most predictive capacity was metabolomics, with R^2^ = 0.176 and P<0.0001 (Figure 6F & G). *Post-mortem* brain volumes displayed the second-best prediction capacity, with R^2^ = 0.095; while R_2_ also yielded significant results, with R^2^ = 0.069. DynaPort gait measurements (R^2^ = 0.07), polygenic scores (R^2^ = 0.05) and routine blood tests (R^2^ = 0.034) also displayed significant results. Other blood molecular data modalities (transcriptomics, proteomics, cytokines, circulating cell-free DNA, pTau-217 & pTau-181) and psychosocial risk factors were not significant (P>=0.05).

#### Ante-mortem predictors of brain resilience molecular subtypes

Continuing our analysis with *ante mortem* predictors, we extend our analysis of associations and development of predictive models to the task of predicting brain resilience subtypes in living adults.

Regarding blood features, only transcriptomic and metabolomic layers displayed significant results (FDR-adjusted P<0.05) in at least one pairwise comparison via permutation ANOVA, controlling for age of death, sex and years of education (Figure 7A). Subtype 1 had 2 upregulated and 10 downregulated metabolomic features when compared to background. For subtype 2, only 1 was significantly upregulated, while 2 were significantly downregulated. For blood transcriptomics, only the comparison between subtype 1 and background yielded significantly differentially expressed features, with 223 transcriptomic features upregulated and 7 downregulated. None of the blood omic layers showed differentially expressed features between subtypes 1 and 2.

**Figure 7.**
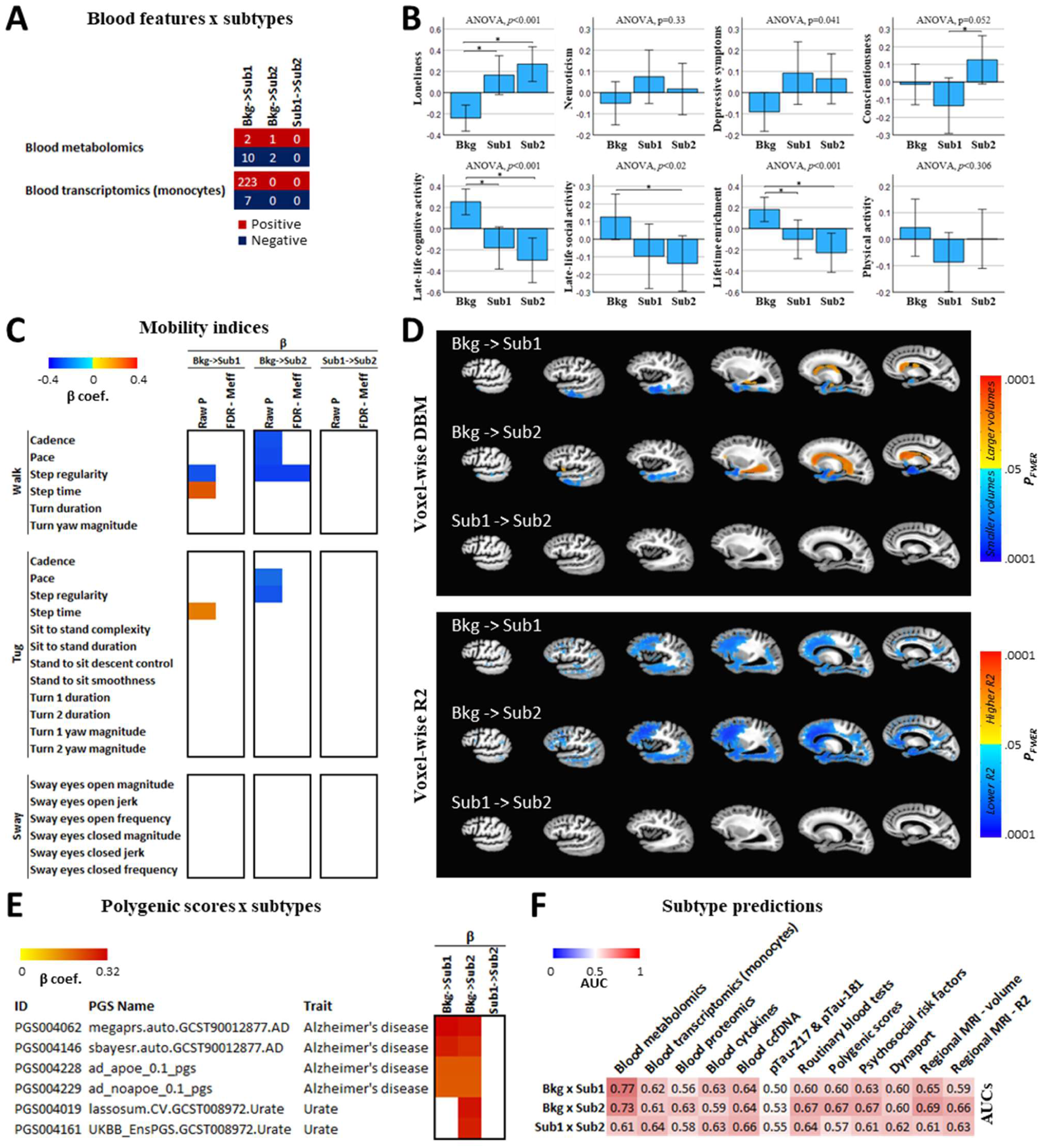
*Ante-mortem* predictions of *post-mortem* brain-derived molecular resilience subtypes. A) Differentially expressed features between subtypes in blood metabolomics and transcriptomics. Red entries represent upregulated features, and blue entries represent downregulated features. Other omic layers not displayed showed no significant results in any comparison. Features were considered significantly differentially altered at FDR-adjusted P<0.05). B) Association of psychosocial risk factors with brain-derived resilience subtypes. Barplots represent z-scored values. ANOVA p-values are displayed above the plots. *: P<0.001. C) Association of gait variables with brain-derived resilience subtypes. Red and blue shaded cells represent, respectively, positive and negative beta coefficients significantly different between each pairwise comparison. Both raw p-values and FDR-adjusted p-values using M_eff_ are presented (both considered significant at P<0.05). Blank cells had non-significant associations (FDR-adjusted P>=0.05). D) Characterization of *post-mortem* brain structure across resilience subtypes. Warm and cold colors indicate the magnitude of positive and negative associations of regional brain deformation (i.e., volumes) and R_2_ (i.e., tissue integrity), respectively (see Methods – Neuroimaging analysis). E) Associations of polygenic scores with brain-derived resilience subtypes. Red shaded cells represent the magnitude of the positive beta coefficients between each pairwise comparison. Only polygenic scores significantly (FDR-adjusted P<0.05) associated in at least one pairwise comparison are shown. Blank cells had non-significant associations (FDR-adjusted P>=0.05). F) *Ante-mortem* cross-validated predictions of brain-derived molecular resilience subtypes. Red shade intensity and numbers inside cells represent AUC values of prediction tasks.

Subtype associations with psychological traits were tested with ANOVA (Figure 7B). ANOVA showed a main effect of resilience subtypes on loneliness (P<0.001), with both subtypes 1 and 2 reporting higher loneliness than the background. We found a borderline main effect of the resilience subtypes on conscientiousness (P=0.052), with *post hoc* comparisons showing that relative to subtype 2, subtype 1 had lower conscientiousness. An overall effect of the resilience subtypes was observed for chronic depressive symptoms (P=0.041); however, we did not find any inter-subtype differences. Likewise, no inter-subtype differences were observed for neuroticism (P=0.33).

Similarly, we tested the associations of lifestyle risk factors with resilience subtypes (Figure 7B). We found a main effect of the resilience subtypes on late-life cognitive activities (P<0.001), with both subtypes 1 and 2 having lower means relative to background. We also found a significant main effect of the subtypes on engagement in social activities (P=0.02), with subtype 2 scoring lower than background. For lifetime cognitive enrichment, we also found a main effect of the resilience subtypes (P<0.001), with both subtypes 1 and 2 having lower lifetime cognitive enrichment relative to background. No differences between the subtypes were observed. We did not observe any subtype differences for engagement in physical activity (P=0.306).

Subtype associations with DynaPort gait measurements were consistent with those observed for pseudotime, with comparison between both resilience subtypes and background samples indicating slower locomotion, with decreases in variables related to speed, and increase in step time (Figure 7C). However, no gait measurement was found to be differentially associated between both subtypes.

Neuroimaging voxel-wise DBM analysis showed that both subtype 1 and 2 displayed smaller temporal regions and larger ventricles compared to background samples (FWER-adjusted P<0.05). However, no significant differences were detected between subtypes 1 and 2 (Figure 7D – upper panel). Additionally, voxel-wise R2 analysis showed lower R2 across brain regions for both subtype 1 and 2 compared to background (FWER-adjusted P<0.05). Again, no differences could be detected between both subtypes (Figure 7D – lower panel).

Polygenic scores also showed associations with subtypes, with 6 PGS displaying positive associations in at least one of the pairwise comparisons (Figure 7E). All 6 PGS were previously associated with pseudotime (Figure 6E). This includes all previously observed AD associated PGS (PGS004062, PGS004146, PGS004228, PGS004229), and two of the PGS associate with urate levels (PGS004019, PGS004161). Urate-associated PGS were positively associated only in subtype 2 compared to background. No PGS was associated in the comparison between both subtypes.

Next, we evaluated the capacity of *ante-mortem* data modalities to accurately predict brain-derived subtypes. We used several machine-learning classifiers with 5-fold cross-validations (see Methods – Statistical analyses). Given the small number of samples with both brain-derived subtype classifications and *ante mortem* data, we conducted pairwise comparisons, trying to analyze each subtype versus background, as well as trying to distinguish between both subtypes.

Results showed great variation in the ability of different data modalities to predict subtype classification (Figure 7F). Metabolomics again was the most promising data layer for subtype prediction, achieving AUC (receiver operating characteristic area under the curve) values of 0.77 when trying to distinguish subtype 1 from background, and 0.73 when trying to distinguish subtype 2 from background. When trying to distinguish both subtypes, however, the prediction power was moderate, with an AUC of 0.61. Circulating cell-free DNA achieved the best discriminative value between subtypes 1 and 2 (AUC of 0.66), although still moderate. Most other data modalities achieved low to moderate values in all comparisons, with AUCs bellow 0.7. Classifications with ptau217 and ptau181 performed worst, displaying essentially no discriminative power (AUCs from 0.50 to 0.55).

Overall, our results show that some different associations can be detected when both subtypes are compared against background samples. However, when trying to distinguish between subtypes, fewer differences can be detected. Regarding our predictive models, while metabolomics achieved the highest predictive power among all data modalities, our models need improvement to achieve a high predictive accuracy. Nonetheless, our results already show the potential for metabolomic data as an *ante-mortem* predictor of brain pseudotime and subtypes.

Given that our model was developed with a limited sample size of 338 individuals, which included those for which we had blood metabolomic data and that were included in the mcTI analysis, substantial improvement in predictive capacity might be achieved if the model is trained using an increased sample size. Additionally, more work can be done to achieve optimal feature selection.

## DISCUSSION

Our work contributes to the characterization of the molecular underpinnings of cognitive resilience, identifying key markers associated with individual variations in resilience and molecular signatures that define heterogenous subtypes of resilience-associated molecular alterations. Importantly, we present the first cognitive resilience pseudo-temporal analysis to our knowledge, while also laying the groundwork for the development of predictive models with the capacity to translate brain-derived molecular pseudotime and subtype classifications to living humans. These represent a first step in enabling the development of novel resilience therapies to maintain cognition in aging adults regardless of which combination of ADRD pathologies is present in the brain.

While previous work trying to characterize the multi-omic molecular signatures of cognitive resilience has been done by our group (*14–18*) and others (*19*), developing an operational metric of resilience capable of quantifying resilience in different individuals has been a challenge. Here, we apply contrastive trajectory inference to develop a quantitative metric of resilience and characterize the multi-omic molecular signatures associated with it.

Previous work from our group using contrastive trajectory inference has also been done in the context of AD with bulk RNA-Seq data from postmortem brain tissues (*24*). This work was expanded by incorporating additional layers of omics data from the same individuals, increasing the amount of molecular information used and enabling us to identify more accurate and meaningful pseudotime values and subtype classifications (*25, 32*). Here, we extend that work to a quantitative metric of resilience. The contrastive trajectory inference has several advantages over quantification based purely on molecular features associated with resilience. First, by building upon contrastive PCA (*43*), contrastive trajectory inference can detect more fine-grained molecular signals associated with resilience by removing common covariance structures between background and target datasets, retaining exclusively those data structures enriched in the target cohort. This approach intrinsically accounts for unknown uninteresting covariables (such as batch effects, demographics, and other pathologies or phenotypes) without having to explicitly control for them. Second, it allows us to calculate a more precise quantification of resilience based on these fine-grained differences identified by contrastive PCA. Third, it can account for the molecular heterogeneity of resilience by identifying distinct subtypes. And lastly, it can accommodate integration of information from more than one omic layer.

Our estimated pseudotime values show strong correlations with cognitive performance (Figure 2B), including residual cognitive performance as expected. The top contributor, *H2AFX*, a variant of histone *H2A*, has a role in DNA damage response (*31*). Phosphorylation of *H2AFX* occurs as a response to DNA double-strand breaks, leading to a relaxation of the chromatin structure around the site of the DNA double-strand breaks and the assembly of the DNA repair machinery at the site. Double-strand breaks are a specially complicated type of DNA damage. Its repair is carried mainly by two pathways: homologous recombination and non-homologous end joining. While homologous recombination can lead to more accurate repairs, non-homologous end joining is more mutagenic. Unrepaired double-strand breaks often leads to apoptosis of damaged cells (*44*). Double-strand breaks accumulate with aging (*45*), and are found to be increased in the brains of patients with mild cognitive impairment and AD (*46*). It has also been suggested that double-strand breaks may have an adaptive role in hippocampus-dependent learning and memory (*47*).

The second top contributor, *UMODL1-AS1*, is a long non-coding RNA with an unknown function. The third, *DXO*, is a decapping enzyme responsible for removing the NAD^+^ (nicotinamide adenine dinucleotide) caps from 5’-end NAD-capped RNAs (*48*). NAD-caps promote RNA decay, contrary to regular 5’-end N7 methyl guanosine (m^7^G) caps, which promote translation and stability. Pseudotime was also shown to be associated with mitochondria-related transcripts and proteins in brain, with a lower expression being associated with lower resilience (higher pseudotime), in line with previous evidence linking mitochondria and bioenergetic capacity to cognitive resilience (*49, 50*). Mitochondrial dysfunction has also been linked to AD (*51*). Additionaly, the only blood omic that had features significantly associated with brain-derived pseudotime was metabolomics, with most metabolites also being downregulated with lower resilience (higher pseudotime). Given the central role that mitochondria have in metabolism, both results seem concordant.

By identifying distinct subtrajectories of pseudotime progression, we decomposed cognitive resilience into two distinct subtypes represented by different molecular signatures, demonstrating heterogeneity in resilience mechanisms. Our two subtypes showed different patterns of cognitive decline and neuropathologies, with subtype 2 displaying an overall less severe profile of decline of several cognitive abilities, as well as a lower burden of AD pathologic traits, but a more severe profile in other ADRD pathologies. Distinct subtypes have been shown before for AD, including molecular subtypes (*52–57*), which highlights the importance of molecular diagnostics in order to reveal heterogenous subtypes of diseases and better inform on potential resilience treatments to maintain cognition and brain health in aging adults.

Another important feature of our study is that we provide our initial efforts to translate the distinct molecular signatures for resilience derived from brain omics obtained from decedents into living adults. Since brain omics cannot be obtained in living adults, we leveraged molecular and clinical data collected from ROSMAP decedents prior to their death to develop predictive models that displayed a promising capacity to estimate pseudotime values and subtype classifications from *ante mortem* data from the same individuals. The most promising *ante mortem* data modality for these predictive tasks was metabolomics, in line with previous findings that showed an association between mitochondria and metabolism with cognitive resilience (*9, 19, 49, 58*).

The study has several limitations. First, the overlap of brain omics is suboptimal. Second the overlap of some ante-mortem datasets is suboptimal. For example, only 185 had Dynaport with post-mortem omics. The numbers with *ante-mortem* MRI were too low for us to use it at all, forcing us to resort to *post-mortem* MRI as a proxy. These issues resulted in a limited capacity to build multi-level classifiers from the clinical data. It is clear from the modest associations of individual features, multiple streams of data will be needed to build multi-level classifiers.

However, this also indicates the potential for substantial improvement of these models with increased sample sizes. Importantly, ROSMAP is currently the only study with as much overlapping data from the same individuals that include survey, clinical, sensor metrics, blood and imaging data paired with multi-omic brain data, making it an ideal source of data for further development of more robust predictive models. The ongoing collection of these data will be crucial for eventually translating brain molecular measures to living persons.

## MATERIALS AND METHODS

### Data

#### Ethics statement

ROS and MAP were each approved by an Institutional Review Board of Rush University Medical Center. All participants enrolled without known dementia and signed an informed consent and Anatomical Gift Act agreeing to annual detailed clinical evaluation and brain donation; in addition, they signed a repository consent allowing their data to be shared.

Data documentation and sharing documents can be obtained at https://www.radc.rush.edu. Study subjects gave written informed consent at the time of enrollment for sample collection and completed questionnaires approved by an Institutional Review Board of Rush University Medical Center.

### Data origin

This study used brain and blood multi-omics molecular, neuropathological, and/or clinical data from the Religious Orders Study and the Rush Memory and Aging Project Study (ROSMAP) (*59*) cohorts.

### Cognitive assessments

Annual administration of 19 cognitive tests were incorporated into a summary measure of global cognition by averaging the z-scores of all 19 tests (*60–62*). For each cognitive measure, the person-specific random slope was estimated as the rate of change in the variable over time, extracted from a linear mixed effects model with the annual variable as the longitudinal outcome, controlling for age at baseline, sex, and years of education (*62, 63*).

### Psychological risk factors

Chronic depressive symptoms were assessed annually using a 10- item form (*64*) of the Center of Epidemiologic Study-Depression Scale (CES-D) (*65*). Chronic loneliness was assessed annually in MAP participants only, using a short version the de Jong-Gierveld Loneliness scale (*66, 67*). We used each participant’s average scores across all evaluations to assess the average level of depressive symptoms and loneliness (*66–69*).

Neuroticism and conscientiousness were assessed at baseline using a short version of the NEO Five-Factor Inventory (*70*). See Table S2 for *ante mortem* data sample sizes.

### Lifestyle risk factors

Physical activity was assessed using questions adapted (*71*) from the 1985 National Health Interview Survey (*72*). Late-life cognitive activity, late-life social activity and lifetime enrichment were assessed in MAP participants only. Late-life cognitive activity was measured with a 9-item scale at baseline (*73*). Late-life social activity was measured at baseline using a 6-item survey on a 5-point scale as reported previously; the mean item score was used in analyses with higher scores indicating greater social activity (*74*). Lifetime enrichment is a composite measure of the mean of early, mid-, and late-life enrichment assessed at baseline.

Early, mid- and late- life enrichment are separate composites. We previously described these composites in detail (*75*). See Table S2 for *ante mortem* data sample sizes.

### DynaPort gait measurements

Annual gait testing session was recorded by a belt-worn sensor positioned over the lower back. This device contains a triaxial accelerometer and gyroscopic to record acceleration and rotation of the lower trunk along each of three orthogonal directions and angular acceleration in three directions. In-home mobility testing recorded four gait tests including: A) a 32ft walk on a 8ft walkway requiring four 8ft walks and three turns; B) two trials of a modified Get Up and Go test (TUG) for 8ft with instructions to stand up from a chair (i.e., sit to stand transition) and walk 8 feet at their preferred pace (i.e., first 8-ft walk), return to the chair (i.e., mid-trial turn, second 8-ft walk), and sit (i.e., stand-to-sit transition), and C&D) Two trials of quiet standing for 20 seconds first with eyes opened and the a second time with eyes closed. Each continuous recording was segmented into individual performances.

Quantitative measures were extracted from each segment. After transformation when appropriate, we used principal component analyses to reduce these measures into gait scores for several of the subtasks, similar to our prior work (*33–35*). See Table S2 for *ante mortem* data sample sizes.

### Polygenic scores

We utilized the PGS Catalog Calculator (*76*) v2.0.0 to calculate polygenic scores (PGS) for ROSMAP individuals with genotyping data. The automated pipeline obtains the scoring files from the PGS Catalog API and applies *liftover* if necessary. We set the genome build to GRCh38, and a minimum percentage of variant overlap was set to 50%. Additionally, no multi-allelic or ambiguous matching variants were allowed. PGS were normalized by ancestry using the HDGP and 1kGP as reference panels. As a result, 802 pre-calculated PGS covering 244 traits were obtained (see File S4 for PGS IDs, names and traits, and Table S2 for *ante mortem* data sample sizes).

### Ante mortem molecular data

The *ante-mortem* blood molecular data was generated in previous studies (*77–79*), and included cell-free DNA (6 features), monocytes RNA-seq (52,650 features; syn22024496), SomaLogic proteins (7,289 features), p-tau181 and p-tau217, cytokines (4 features), and/or Biocrates metabolites (292 features) for a subset of 1,108 ROSMAP participants with brain multi-omics data (see Table S2 for *ante-mortem* multi-omic data sample sizes).

### Post-mortem imaging data

After one month postmortem, the cerebral hemisphere pre-selected for neuropathological examination was imaged with a turbo spin echo sequence on one of four 3- Tesla scanners, as previously reported (*80*). We generated deformation and R_2,_ the inverse of T_2_, maps as previously reported (*81–84*). To generate features for machine learning analyses, we also generated regional grey matter estimates using an in-house multi-atlas segmentation pipeline transformed into subject space. More details on this pipeline may be found elsewhere (*85*).

### Neuropathologic data

All subjects underwent *post mortem* neuropathologic evaluations, including uniform structured assessment of AD pathology, cerebral infarcts, Lewy body disease, TDP-43 cytoplasmatic inclusions in neurons and glia, cerebral amyloid angiopathy, hippocampal sclerosis, and atherosclerosis (*12, 86*).

### Post-mortem omics data

The *post-mortem* brain multi-omics data included bulk RNA-seq expression and tandem mass tags (TMT) proteomics from the dorsolateral prefrontal cortex (DLPFC) of 898 autopsied brains (see Table S1 for demographic characteristics and Table S2 for multi-omic data sample sizes), generated in previous studies (*18, 87, 88*). Participants were included if they had at least one of the two data modalities, and a resilience cluster classification (see Methods – Clustering subgroups of residual cognitive decline). Most of these data is available at the Accelerating Medicines Partnership Alzheimer’s Disease knowledge portal (AMP-AD; www.synapse.org). The following Synapse IDs were used: syn3388564 and syn17015098. Bulk RNA-seq (syn3388564) was extracted using the Chemagic RNA tissue kit and sequenced with NovaSeq 6000 (Illumina) (*88*). Proteomic data (syn17015098) was generated for a total of 8,391 protein levels quantified using tandem mass tags (TMT) (*18*).

### Data preprocessing

Before analysis with the *mcTI* algorithm, transcriptomic and proteomic feature’s values were adjusted for age at death, sex and years of education using robust additive linear models.

### Clustering subgroups of residual cognitive decline

We used a latent class functional mixed-effects model (*26*) to identify clusters of individuals who displayed similar patterns of residual cognitive decline. We included individuals: (i) without dementia at baseline; (ii) with at least one follow-up; (iii) deceased with brain autopsy and a completed neuropathologic exam; and (iv) without other major pathological diagnoses, e.g., large brain tumor. In total, 1409 individuals were selected for clustering. After controlling for demographic (age at death, sex and years of education) and neuropathologic indices (AD, TDP- 43, cerebral amyloid angiopathy, Lewy bodies, hippocampal sclerosis, atherosclerosis, macroinfarcts and microinfarcts), the functional mixed effects model assumes that the random coefficients follow a mixture distribution with *G* latent classes, where the optimal number of latent classes *G* was determined by minimizing the Bayesian information criterion. Participants were subsequently classified into corresponding latent classes based on posterior probabilities estimated from the model. Four resilience clusters were identified, representing four distinct trajectories of residual cognitive decline. For mcTI analysis, resilience cluster 1, individuals with no residual cognitive decline, was used as the background; and resilience clusters 3 and 4, those with the steepest residual cognitive decline, were combined and used as the target, as cluster 4 alone had two few persons for stable modeling.

### Multimodal contrastive Trajectories Inference (mcTI)

Initially proposed in (*25*), the mcTI algorithm has been updated to improve specific aspects (*32*). Given a set of multiple ‘omic’ data types, this method provides two estimates for each participant: 1) molecular disease pseudotime reflecting how close (in terms of multi-level molecular alterations) each participant is to developing the condition of interest (i.e., faster residual cognitive decline), and 2) a subtype which corresponds to a distinct subtrajectory of pseudotime progression from the healthy background state to the most severe state. Before the analysis, the high dimensionality of each data modality is reduced by selecting the 1000 features with the highest variance in the target population (resilience clusters 3 and 4).

First, a contrastive Principal Component Analysis (cPCA) (*43*) is performed, reducing each data modality to a few components capturing the resilience clusters 3 and 4 associated patterns (*24*). The contrasted principal components (cPC) resulting from all different data modalities are subsequently used as input in a probabilistic Principal Component Analysis (pPCA) which identifies fused contrasted components capturing the common variance across all modalities’ contrasted/enriched patterns, while dealing with missing data, i.e., varying N’s for each omic layer. A subject-subject dissimilarity network is then constructed based on the inter-individual Euclidian distances across the resulting fused cPCs. The dissimilarity network is used to calculate the shortest path from any participant to the background subjects. Each shortest path is defined as the concatenation of relatively similar subjects in the integrated multi-omics molecular space that minimizes the distance to the background’s centroid. Relatively low or high values indicate greater or lesser distance on the path to the fastest residual cognitive decline clusters 3 and 4.

Once the pseudotimes are estimated, before proceeding to identify subtypes, all the fused contrasted components are statistically adjusted by the pseudotime values via robust regression. These new disease-adjusted components are then non-linearly embedded into a two- or three-dimensional space via t-SNE (*27*), where the participants are subsequently clustered. The number of t-SNE dimensions is selected to maximize results stability across perplexity values (i.e., in the range [5 to 50] with 5 as step size) and clustering criteria. The resulting clusters define the final putative subtypes. To identify the optimum number of clusters/subtypes, a majority rule across the Calinski–Harabasz and Silhouette criteria is used (*28, 29*). The subtypes’ stability and significance are evaluated via randomized permutations. Following the method proposed in (*52*), subtype stability is defined as the rate at which pairs of subjects group together into the same subtypes upon repeated clustering on random subsets of the input data. Analyses were performed in MATLAB version R2021b.

### Assessing features’ contributions to pseudotime

For each dataset and molecular omics modality, the total contribution 𝐶_i_ of each modality-specific feature *i* to the obtained reduced representation space (and the multi-omics pseudotime) was quantified as (*25*):

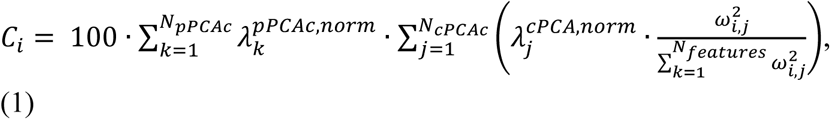

where 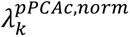 is the normalized eigenvalue of the pPCA component *k* from a total of 𝑁_pPCAc_ resulting components after fusing all data modalities, 𝑁_cPCAc_ is the number of contrasted principal components for marker *i*’s corresponding data modality, 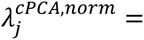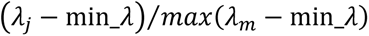 is the normalized eigenvalue of the contrasted principal component *j*, min_𝜆 is the minimum obtained eigenvalue across all markers for *j*, 𝜔_i,j_ is the loading/weight of the marker *i* on the component *j*, and 𝑁_markers_ is the total number of modality-specific markers in *i*’s corresponding data modality.

For comparisons across data modalities, the contribution values were normalized by the value of the highest contributor across the modalities.

### Identification of distinct subtype-specific signatures

We identified distinct molecular alterations across each subtype by detecting omic features that deviated from typical values in the resilience cluster 1 reference group (P<0.05, FDR-corrected), while remaining stable in other subtypes. For each subtype *i* and omic feature *j*, we employed an ANCOVA test with 2,000 randomized permutations to calculate their differentially altered levels, generating (F*_i,j_* and P*_i,j_*) statistics, adjusted for age at death, sex, and years of education. We then identified all omic features that were significantly different for any subtype (P<0.05, FDR-corrected). From this pool, we selected the omic features that were altered for subtype *i* but not altered for the other subtypes. These selected omic features constituted the distinct molecular signature for subtype *i*.

For interpretation, within subtype *i*, the selected features were also ranked according to their propensity to be predominantly different for *i* while being preserved in the other subtypes. To quantify this tendency, we calculated an ’anisotropy’ index, defined as follows (*89, 90*):

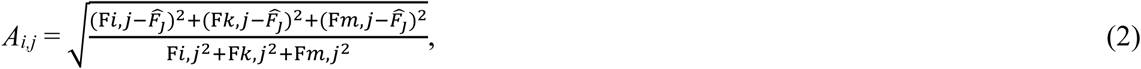

where *k* and *m* represent the other subtypes and 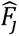 is the average F-value for feature *j* across subtypes. This formula can accommodate different numbers of subtypes by including or excluding terms accordingly. The index (*A_i,j_*) yields a scalar value between zero and one, indicating the distinctiveness of feature *j* concerning subtype *i*; values closer to one suggest a stronger distinctiveness. This approach is an extension of the concept of fractional anisotropy commonly employed in diffusion neuroimaging and relates to the eccentricity of conic sections in three dimensions, normalized to a unit range. Notice that while F-values may not be directly comparable across data modalities due to varying sample sizes, the *A_i,j_* values allow for meaningful comparisons since they are standardized across subtypes.

### Gene Ontology (GO) enrichment analysis

Features were enriched for GO terms (version 2025-03-16) (*91*) using the *rbioapi* R package v0.8.2 (*92*). All features in our initial dataset were used as the background set. P-values were FDR-adjusted and considered significant at adjusted-P<0.05. Only overrepresented terms were considered.

### Neuroimaging analysis

To examine the voxel-wise associations of molecular pseudotime and resilience subtypes with brain morphology and tissue integrity, we implemented two sets of general linear models (GLMs) using FSL PALM (*93, 94*), which could assume different variances across scanners. In the first model, we entered resilience pseudotime as the explanatory variable (EV) of either regional deformation or R_2_ and controlled for age at death, sex, years of education, postmortem interval, and scanner location. In the second model, we entered three indicator variables for the background and resilience subtypes as EVs of interest, also adjusted for demographics and postmortem conditions, and ran contrasts for each group pair. P-values were computed from 500 permutations using tail approximation (*95*), threshold-free cluster enhancement (TFCE), and family-wise error rate (FWER) correction. Associations with FWER *p*≤.05 were considered statistically significant. These analyses were performed using 536 decedents with *post-mortem* neuroimaging (Bkg = 237, Sub1 = 124, and Sub2 = 175). See Table S2 for *ante mortem* data sample sizes.

### Statistical analyses

Associations of brain-derived resilience pseudotimes with cognitive, neuropathological and *ante- mortem* variables were estimated via linear regression. Associations with resilience subtypes were estimated via ANOVA with 2,000 permutations for continuous variables and Chi-squared test for categorical variables. Strength of association for Chi-squared tests were determined using Cramer’s V. Given that molecular pseudotime and subtypes were already derived from an algorithm controlling for age at death, sex and years of education, analyses did not further control for these covariables, unless stated otherwise. Machine-learning prediction tasks with *ante mortem* data of resilience pseudotimes used linear regression, SVM, classification trees, ensembles, neural networks, and gaussian process regression, under a 5-fold cross-validation scheme. For resilience subtypes, classifications with *ante mortem* data used SVM, classification trees, naïve Bayes, neural networks, *k*-nearest neighbor, linear discriminative analysis and ensembles, under a 5-fold cross-validation scheme. Prediction performance metrics (R^2^ and AUC) reported in this paper represent the best predictions across all methods tested. All prediction tasks included age at death, sex and year of education as covariables.

## Supporting information

File S1 - Regression results of pseudotime association with features

File S2 - GO enrichment of top features associated with pseudotime

File S3 - Distinctive subtype signatures

File S4 - Polygenic scores

Table S1 - Demographics of participants

Table S2 - Post mortem and ante mortem data sample sizes

## Supplementary Materials

**Table S1 |** Demographics of participants.

**Table S2 |** Post mortem and ante mortem data sample sizes.

**File S1 |** Regression results of pseudotime association with features.

**File S2 |** GO enrichment of top features associated with pseudotime.

**File S3 |** Distinct subtype signatures.

**File S4** | Polygenic scores.

## Data and materials availability

ROSMAP omics data are available for general research at the AMP-AD knowledge portal (https://www.synapse.org/), according to the following requirements for data access and attribution (https://adknowledgeportal.synapse.org/DataAccess/Instructions). ROSMAP neuropathological, neuroimaging, psychological and clinical data are available at the RADC Research Resource Sharing Hub (<u>radc.rush.edu</u>), pending scientific review and a completed material transfer agreement (see <u>radc.rush.edu/requests.htm</u>). mcTI is freely available as part of the *NeuroPM-box* software (*96*) (<u>neuropm-lab.com/neuropm-box.html</u>).

## Acknowledgments

We appreciate the participants of ROS and MAP for their generous gift and thank the staff of Rush Alzheimer’s Disease Center for data collection, management, and analyses. The results published here are in whole or in part based on data obtained from the AD Knowledge Portal (https://adknowledgeportal.org ). Study data were provided by the Rush Alzheimer’s Disease Center, Rush University Medical Center, Chicago. Data collection was supported through funding by NIA grants P30AG10161, P30AG72975 (ROS), R01AG15819 (ROSMAP; genomics and RNAseq), R01AG17917 (MAP), R01AG36836 (RNAseq), R01AG48015 (monocyte RNAseq), U01AG32984 (genomic and whole exome sequencing), U01AG46152 (ROSMAP AMP-AD, targeted proteomics), U01AG46161, U01AG072572 and U01AG061357 (TMT proteomics), U01AG61356 (whole genome sequencing, targeted proteomics, ROSMAP AMP-AD), R01AG59732, R01AG75728, R01AG79133, P30AG72975, U01AG046152, R01AG30146, the Illinois Department of Public Health (ROSMAP), and the Translational Genomics Research Institute (genomic). Additional phenotypic data can be requested at www.radc.rush.edu. YIM is supported by a Canada Research Chair tier-2, CIHR Project Grant 2020, and Weston Family Foundation’s Transformational Research in AD 2020. TWC is supported by The Phil and Penny Knight Initiative for Brain Resilience. PA is supported by the National Institute on Aging: P30AG021334 (Johns Hopkins Older Americans Independence Center), K24AG088484, P30AG073104 (Artificial Intelligence and Technology Collaboratory for Aging Research). We would also like to thank Michael Urbut and the Paul M. Angell Family Foundation (SDA-F24-08) for their generous financial support given towards this work. The funding organizations had no role in the design or conduct of the study; collection, management, analysis, or interpretation of the data; or preparation, review, or approval of the manuscript.

