## Supplementary material for "Characterizing Post-Mortem Brain Molecular Taxonomy of Cognitive Resilience and Translating it to Living Humans": Table S1 - Demographics of participants

**Table S1.** Demographics of participants - descriptive characteristics of the whole sample and stratified by background and non-background groups.

| Characteristics | Whole sample | Background | Non-background  Target | Non-background  Other* |
| --- | --- | --- | --- | --- |
| N | 898 | 410 | 239 | 249 |
| Age at death, mean years (SD) | 90.1 (6.4) | 89.1 (6.3) | 91.2 (6.3) | 90.7 (6.5) |
| Female, n (%) | 635 (70.7) | 276 (67.3) | 179 (43.7) | 180 (72.3) |
| Educational attainment, mean years (SD) | 16.1 (3.6) | 16.2 (3.5) | 16.3 (3.9) | 15.9 (3.6) |

*Non-background samples that were not set as target.
