## Supplementary material for "Characterizing Post-Mortem Brain Molecular Taxonomy of Cognitive Resilience and Translating it to Living Humans": Table S2 - Post mortem and ante mortem data sample sizes

| Data modality | | Sample size |
| --- | --- | --- |
| ***Brain post mortem*** | |  |
| RNA-Seq | | 844 |
| TMT proteomics | | 613 |
| ***Matched ante mortem*** | |  |
| Blood transcriptomics | | 212 |
| Blood proteomics | | 715 |
| Blood metabolomics | | 338 |
| Blood cytokines | | 182 |
| Blood circulating cell-free DNA | | 182 |
| Blood p-tau181 and p-tau217 | | 138 |
| Psychosocial assessments | | 256 |
| DynaPort gait measurements | | 185 |
| Polygenic scores | | 875 |
| Neuroimaging | | 535 |

**Table S2.** *Post mortem* and *ante mortem* data sample sizes.
